## Supplementary figures and legends for "Distinct neural mechanisms of social orienting and mentalizing revealed by independent measures of neural and eye movement typicality"

**Supplementary Movie 1: Average, most typical, and least typical eye movements.** One of the 24 movie clips, with the mean scan path of all the TD participants depicted by the magenta circle, fixations of the most typical participant shown by the black circle, and of the least typical participant shown by the white circle. This is the same movie clip portrayed in Figure 1. Movies were presented to participants at a higher resolution (1280*1024) and with the corresponding soundtrack, unlike this clip which was generated post hoc with the overlaid fixations.

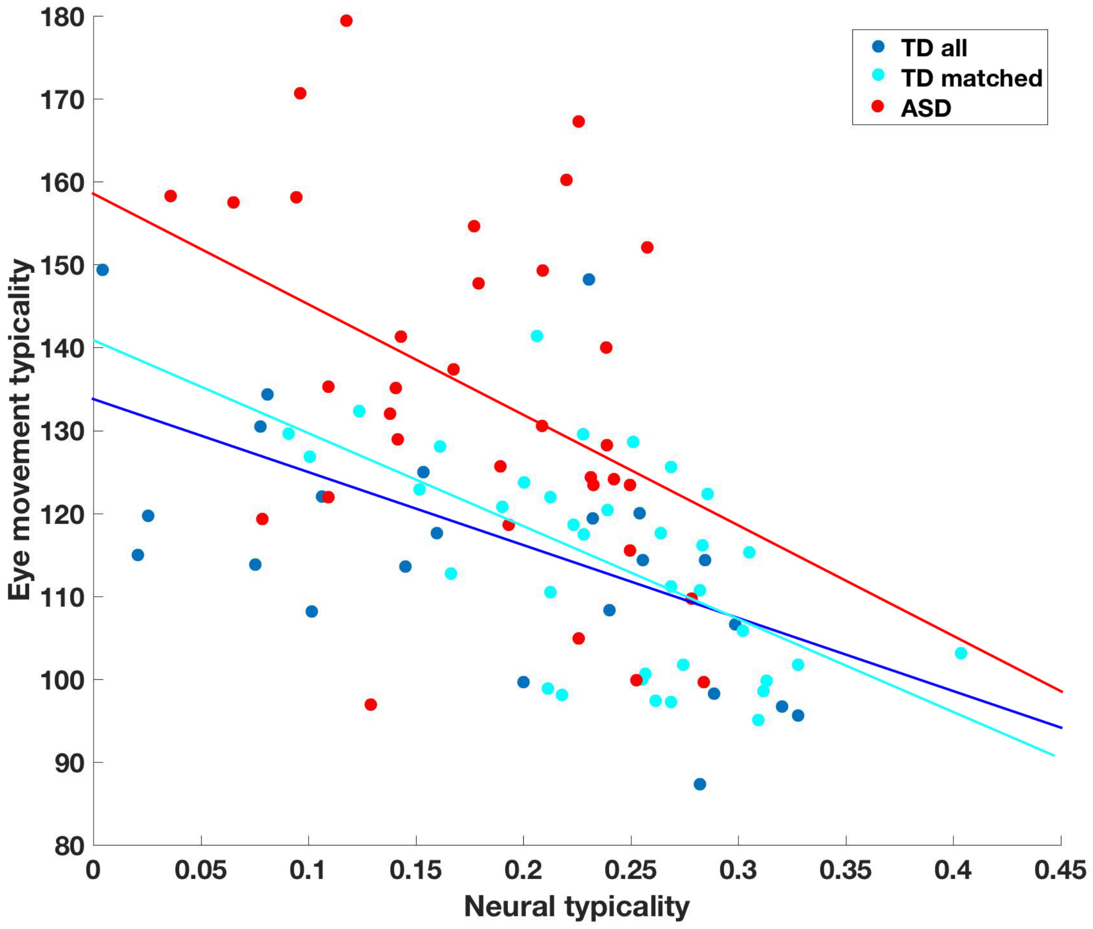


**Supplementary Figure 1: Correlations between neural typicality and eye movement typicality using across group definitions**. Correlation between eye movement typicality and neural typicality for the ASD group (red), averaged across the social orienting network as defined by the correlations of eye movement and neural typicality for the TD group. R = -0.42, p = 0.01. Correlations between the eye movement and neural typicality for the matched TD subset (cyan, r = -0.59, p = $1.6*10^{-4}$), and the entire TD group (remaining participants shown in blue, r = -0.56, p =$4.2*10^{-6)}$, shown for context.

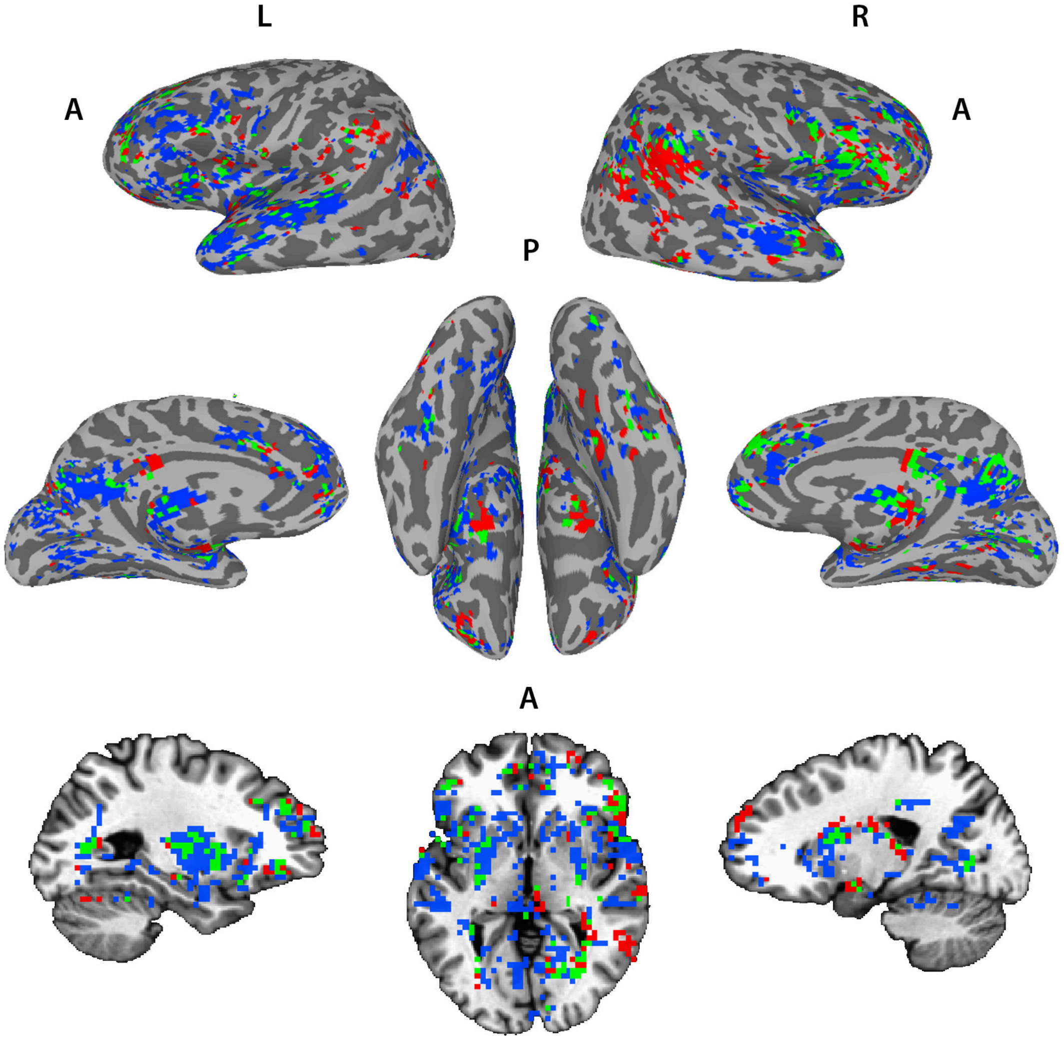


**Supplementary Figure 2: conjunction map of neural typicality group differences and eye movement correlations**. Overlay of the voxels with significant correlations between neural typicality and eye movement typicality of the combined matched TD and ASD groups (N = 36 TD + 33 ASD, blue), voxels showing significant group differences in neural typicality between the matched TD and ASD groups (N1 = 36 TD, N2 = 36 ASD, red), and voxels significant in both analyses (green). Threshold set at p<0.01, corrected for multiple comparisons through cluster size permutation tests.
